# Nanopore-Based Whole-Genome Sequencing Reveals a Bioluminescent and Non-Bioluminescent Bacterium Isolated from the Squid *Loligo forbesi* in the Red Sea

**DOI:** 10.1101/2025.04.27.649427

**Authors:** Mohammed A. Hassan, Ahmed A. Radwan, Abd El-Nasser A. Khattb

## Abstract

Common in aquatic life, symbiotic interactions with luminescent bacteria to enable bioluminescence. Although interactions in cephalopods have been well investigated in species including *Euprymna scolopes*, little is known about other cephalopods, especially those found in the Red Sea. This work separated and identified bioluminescent and non-bioluminescent bacterial strains from the ink sacs and eyes of *Loligo forbesii*, a squid species taken from the Zaafarana area of the Suez Canal in Egypt’s Red Sea. Whole-genome sequencing based on Nanopores turned up two distinct bacterial strains: *Shewanella baltica* (non-bioluminescent) and *Psychrobacter alimentarius* (bioluminescent). Under Alkanal Monooxygenase (FMN-linked), genome annotations in the *P. alimentarius* genome expose Alkanal Monooxygenase Alpha Chain bacterial bioluminescence. High similarity between the isolated *P. alimentarius* strain and publicly archived reference genomes was confirmed by a comparative genomic study, including an ANI calculations tool and phylogenetic tree building. Moreover, a whole-genome alignment using the Mauve tool revealed a high level of sequence conservation between our *P. alimentarius* isolate and previously published strains in GenBank. The findings suggest that *P. alimentarius* may be involved in the bioluminescence of *L. forbesii* and support current understanding of cephalopod bacteria associated with *L. forbesii*. The present work provides more information on the whole genome sequence including genomes annotation of both bioluminescent and non bioluminescent strain and explores the bacterial symbiotic in cephalopod marine organisms inhabiting the Red Sea.

## 1. INTRODUCTION

Including cephalopods, many terrestrial and aquatic species emit light. Moreover, bioluminescence (Haddock *et al*., 2010; Otjacques *et al*., 2023) functions as both a mechanism of attraction and defense. In the generation of light, a photophore is an organ. Although the autogenic organisms create light by themselves, the bacteriogenic organisms create light by the bioluminescent bacteria in a symbiotic relationship with the organism (Haddock *et al*., 2010). Recent studies on bioluminescent cephalopods, such as *Euprymna scolopes*, generated light by means of bioluminescent bacteria *Vibrio fischer*i. *Vibrio fischer*i extracellularly colonizes *E. scolopes* along their apical surface of epithelium. Human microbiome cells interact with gut epithelia (Nyholm and McFall-Ngai, 2021) similarly to this mechanism. By means of the epithelia and the processes by which they survive over the lifetime of the host, the squid-vibrio system can also help one understand the mechanisms behind the recruitment of gram-negative bacteria (Otjacques *et al*., 2023). Combining ventral photophores containing bioluminescent bacteria, the *Uroteuthis* genus of loliginid squids (Cephalopoda: Loliginidae). Several paired structures have been shown to exist in the ink sacs of female *Afrololigo mercatori*s and young *Loligo forbesi* (Alexeyev, 1992). Species of loliginid squid were categorised under subfamily Uroteuthinae (Alexeyev, 1992). These species have photophores, although the accessory nidamental glands of *L. forbesi* (Lum-Kong and Hastings, 1992; Vecchione *et al*., 1998) revealed a luminescent bacterial strain. A species of Psychrobacter (accession number LK9318) has revealed a fresh example of bioluminescence, that is, light emission.

The first known documentation of Bioluminescent within this class of bacteria comes from presence of a symbiotic association with the rani fish (*Nemipterus japonicus*), the bioluminescent *Psychrobacter* is Luminescence remarkably increased as the *Psychrobacter* genera in the culture entered the mid-logarithmic development phase. This phenomenon called quorum sensing (Prakash *et al*., 2014), bacteria coordinate activities using a communication system based on population density. These days, quorum sensing is studied extensively by researchers to investigate related mechanisms in other species as well as the underlying cellular communication channels and biochemistry engaged (Williams *et al.,* 2007). It has been suggested to assist photoreactivate DNA repair systems, neutralise damaging reactive oxygen species, and offer an extra layer of protection against the negative effects of short-wavelength solar ultraviolet (UV) radiation. Additionally involving this light production are mutualistic interactions involving different cell types or to attract them. The supposed mutualistic symbiotic relationship between *Shewanella pealeana* and *Loligo pealeii* is one-sided. Linked to the last stage of egg development, the accessory nidamental gland (Bloodgood, 1977; Leonardo *et al*., 1999) is typified in molluscs by a gland rich in bacteria. Living in squid eggs, *S. pealeana* guards against abiotic stresses, pathogens, predators, and parasites (Barbieri et al., 1997). Comprising the α (LuxA) and β (LuxB) subunits, LuxAB is the particular enzyme complex in charge of this light generation. LuxAB produces light while being connected to a molecule called reduced flavin mononucleotide (FMNH[) and using molecular oxygen to change a long-chain aldehyde into a fatty acid. Research on bioluminescence exposed a suite of bacterial species with rearranged *lux* operons. These species lack a component of luxB. *Escherichia coli* can produce luminescence when introduced by a synthetic *lux*A gene derived from a simpler *lux*ACDE operon found in *Enhygromyxa salina*. Overall, the *Es*LuxA gene generates less light than the luciferase genes seen in *Aliivibrio fischeri* (*Af*LuxAB). Discovered in coastal Japan, the rather halophilic myxobacterium *Enhygromyxa salina* is not known to be luminous. Compared to usual heterodimeric luciferases, *Es*LuxA’s luminosity is far less (Yudenko *et al.,* 2024).

In Japanese coastal, the rather halophilic myxobacterium *Enhygromyxa salina* is not reported to be luminous. *Es*LuxA’s luminosity is considerably less than that of typical heterodimeric luciferases (Yudenko *et al.,* 2024). *Vibrio fischeri* is the most famous bacterial bioluminescence, the emission of light by bacteria, has been extensively investigated in many marine environments and estuaries. Visibly luminous (producing high levels of light), non-visibly luminous (producing low levels of light, detectable only with a photometer), or non-luminescent (producing no light) are the several ways that bioluminescent bacteria might be classified. Fascinatingly, studies have found luciferase gene sequences in even nonluminescent and nonvisibly luminous strains. For instance, the reference strain *V. cholerae* ATCC 145 might generate just low levels of light even though it contains the *lux*A gene required for bioluminescence. This phenomenon implies that the existence of luciferase genes does not always correlate with visible light emission, maybe due to a lack of other genes in the lux pathway, uncoordinated gene control, or changes in other genes involved in the bioluminescence process (Ramaiah *et al*., 2000). By means of Nanopore-based whole-genome sequencing, the present work aims to identify a bioluminescent strain and a non-bioluminescent strain living symbiotically in several regions of the squid’s body.

## 2. MATERIALS AND METHODS

### 2.1. Isolation and Culturing of bioluminescent bacteria

The squid samples, *L. forbesii*, were collected from the Zaafarana area (Suez Canal, Red Sea, Egypt), and kept in an icebox and transported to the Biotechnology Institute of the National Research Centre (NRC) for microbial genetic study. To get rid of sand or mud, the squids were gently cleaned using sterilized distilled water. The squids were then put in seawater, half submerged for several hours at 4°C to 8° refrigerated temperature.

Following dissection of the squid digestive system and ocular structures, ink samples were taken from the ink sacs and eyes of the squids, illustrated in **Figure 1** (b and C). Serial dilutions were carried on until a 10^-5 dilution was obtained. Every dilution went on LA plates (Peptone 10 g/L; MgSO_4_ 1 g/L; K_2_HPO_4_ 4 g/L; NaCl 30 g/L; Glycerol 60%) , modified from that described by Nealson, 1978. Avoiding UV lamp illumination, the plates were incubated at 25°C for 24 hours, with luminescent colonies observed and recorded every 8 hours in a dark chamber of the Gel documentation device. Random selection produced brightest light-emitting colonies; these were streaked onto LA plates and incubated for another 24 hours at around 25°C. After that, the unique luminescent bacterial colonies were identified through luminescence observation and purified by sub-culturing on LA plates using standard streaking techniques. Pure cultures were subsequently stored in LB for further growth and analysis.

**Figure 1:**
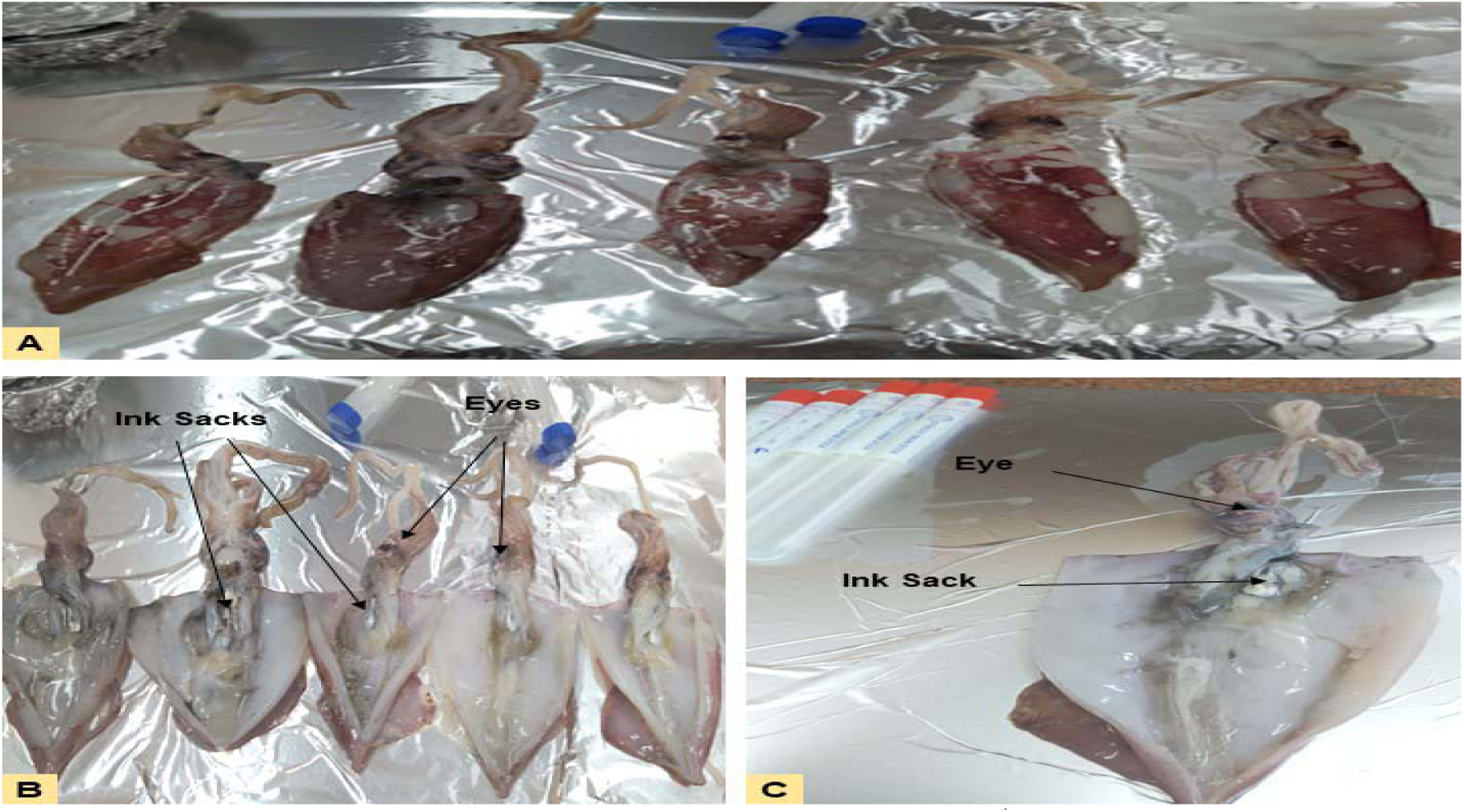
The squid samples that could provide bioluminescent bacteria. **A)** The collected squids, *L. forbesii*, from the Zaafarana region. **B, C)** illustrate the position of a squid’s eye and show its dissection. The black arrows show the particular sites, the eye and the ink sac (silver body) from which sterile swaps were taken.

### 2.2. Genomic DNA Isolation and 16s rDNA PCR Reaction

A modified approach (Williams *et al*., 2012) was used to extract high-molecular-weight (HMW) DNA. Using a NanoDrop microvolume spectrophotometer, DNA concentration was measured; quality was evaluated by loading samples onto a 0.8% agarose gel in 1× TAE buffer to validate integrity and absence of smudging (Zhang *et al*., 2024). Three PCR reactions were optimized to amplify the 16S rDNA template using a melting temperature of 52°C to investigate whether the extracted HMW of genomic DNA degrades the 16s rDNA templates. Using the Rapid Barcoding Kit method, HMW DNA was delivered to a genomic services facility at the Children’s Cancer Hospital for sequencing on the Nanopore platform (MinION Mk1C).

### 2.3. Genome Assembly

MinION readings went through a multi-step quality control process. First FastQC (version 01-03-23), then adapter removal with Porechop (version 0.2.4) to examine the raw data. Further filtered using Filtlong (version 0.2.1), which used k-mer matching to a reference sequence for improved quality assessment, were reads using Flye (version 2.9.5), a de novo assembler idealized for long-read data, filtered reads were reassessed under FastQC before assembly (Lin *et al*., 2016). Flye can tolerate high error rates and process varying sized genomes. Racon (Vaser *et al*., 2017) aligned reads to contigs and corrected mistakes, so improving the assembly. Bandage (Wick *et al*., 2015) visualized assembly graphs; assembly quality was assessed with QUAST (Gurevich *et al*., 2013).

#### 2.3.1. Genome Contamination

Accessible via the Biocloud server, the ContEst16S online tool was used to evaluate the possible contamination risk so guaranteeing the purity and integrity of the genome. This tool identifies contamination by analyzing 16S rRNA gene sequences within the assembly (Lee *et al*., 2017).

#### 2.3.2. Genome Annotation Using DFAST

The finalized assembled genome was annotated and subsequently submitted to the DNA Data Bank of Japan (DDBJ) at the National Institute of Genetics. This was accomplished using the DDBJ Fast Annotation and Submission Tool (DFAST), accessible at https://dfast.ddbj.nig.ac.jp.

#### 2.3.3. Analyzing and Visualizing Genomic Data with BV-BRC

The Bacterial and Viral Bioinformatics Resource Centre (BV-BRC) server (https://www.bv-brc.org/) annotated the whole genome. The circular viewer program proKsee (Grant *et al*., 2023) let only view annotated genomic sequences.

#### 2.3.4. ANI Calculator

OrthoANIu (https://www.ezbiocloud.net/tools/ani) is an improved method grounded on USEARCH that helped to estimate the average nucleotide identity (ANI) between genome sequences. Yoon et al. (2017) claim that this method presents a simpler and more successful substitute for the first developed OrthoANI approach.

#### 2.3.5. Constructing Bacterial Phylogenetic Trees: Genomic and 16S rRNA Sequence Analysis

Using Mash/MinHash for the estimate of genetic similarity, the Similar Genome Finder in BV-BRC was used to identify bacterial genomes closely associated to the query genome (Ondov *et al*., 2016). Using the codon tree approach, which finds single-copy genes from BV-BRC PGFams and uses RAxML for analysis, a phylogenetic tree was built. MAFFT (version 7.526) generated alignments of 16S rDNA sequences that were visualised using Archaeopteryx (https://www.phylosoft.org/archaeopteryx) (Katoh and Standley, 2013).

#### 2.3.6. Genome Alignment (Mauve)

Whole-genome alignment was performed using Progressive Mauve (Darling *et al.,* 2010) to identify conserved regions and genomic differences.

## 3. RESULTS

### 3.1. Cultivation and Selection of the Luminescent Bacteria

The dark settings facilitate selection of the luminescence strain. The bioluminescent bacteria linked to ink sacs and eyeballs were grown on LA plates and imaged in a dark chamber, see **Figure 2**. The colonies that shows up on the solid medium as a white dot under normal light and exhibits luminescence in dark were selected for further investigation.

**Figure 2:**
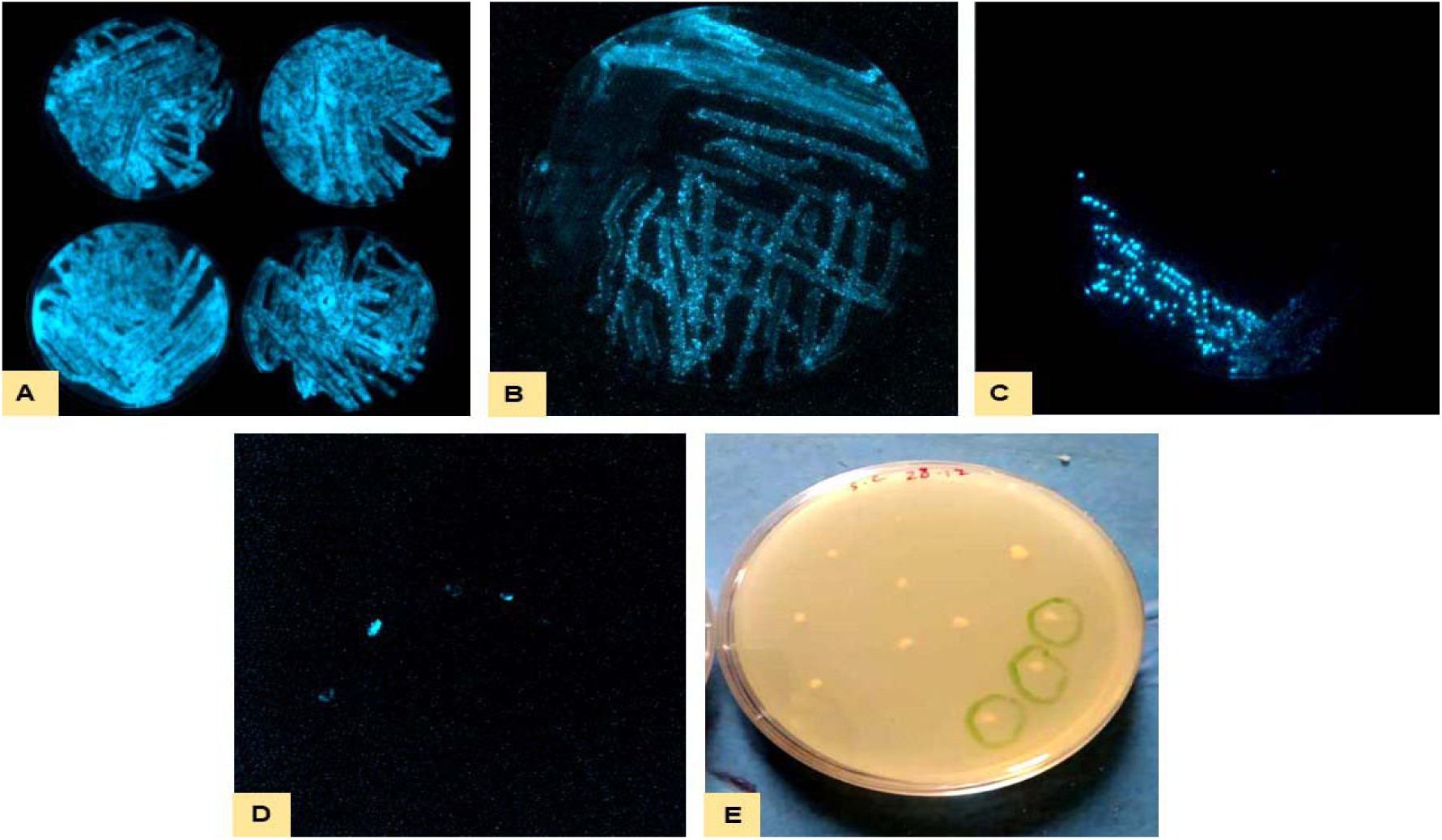
The luminescent bacteria that isolated from the ink sacs and eyeballs of *Loligo forbesii* grown on specialized luminescent agar medium imaged in a dark chamber (**A**, **B**, **C**, and **D**) and under normal light (**E**).

### 3.2. High Molecular Weight DNA Isolation and Detection of 16s rDNA

We isolated high molecular weight (HMW) DNA from the luminescent broth of pure cultures. The extracted DNA concentration was 300 ng/µl as measured by NanoDrop. After extraction, yielding the complete genome was confirmed through three PCR reactions for 16S rRNA region. A sharp band corresponding to the 16S rDNA gene was visualized on an agarose gel with an expected molecular weight of 1600 bp, as shown in **Figure 3**.

**Figure 3:**
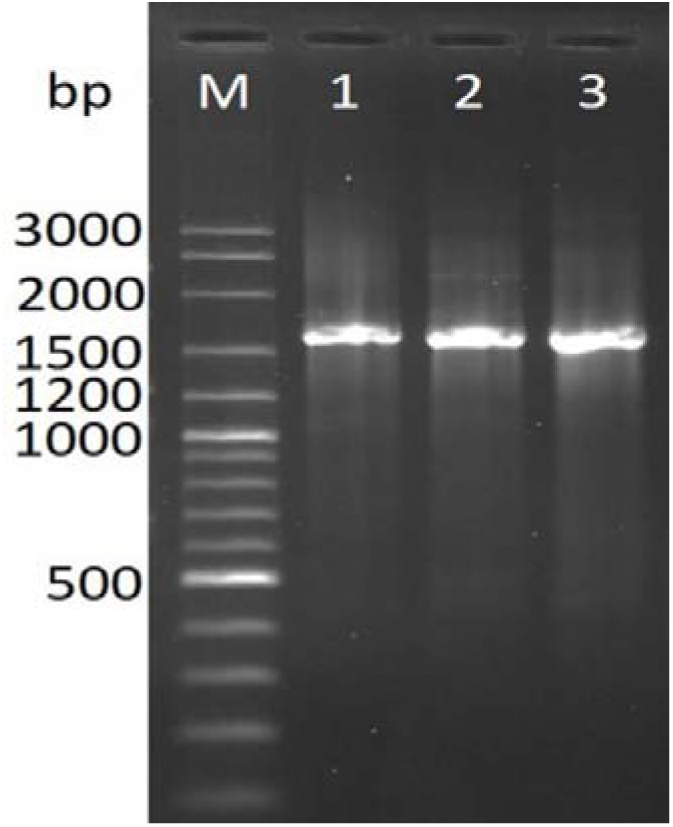
Detection of PCR products of 16S rDNA region at ≈ 1600 bp.

### 3.3. *De novo* Assembly and Quality Control Results

Following confirmation that the 16S rDNA PCR product was good, we generated high molecular weight (HMW) genomic DNA using the Nanopore Rapid Barcoding Kit and then put it onto a MinION Mk1C flowcell. Fastq files were then handled according to the assembly processes and quality control guidelines found in the Materials and Methods section. We handled the Fastq files according to the assembly and quality control procedures described in the Materials and Methods section. Generating assembly graphs and summary reports, we validated the final genomic assembly using the Bandage tool. As seen in **Figure 4**, two main nodes were found: the smallest contig (3,256,386 bp) and the largest contig (4,992,825 bp). Reduced numbers of contigs, according to Quast, point to a better-quality assembly with less mistakes.

**Figure 4:**
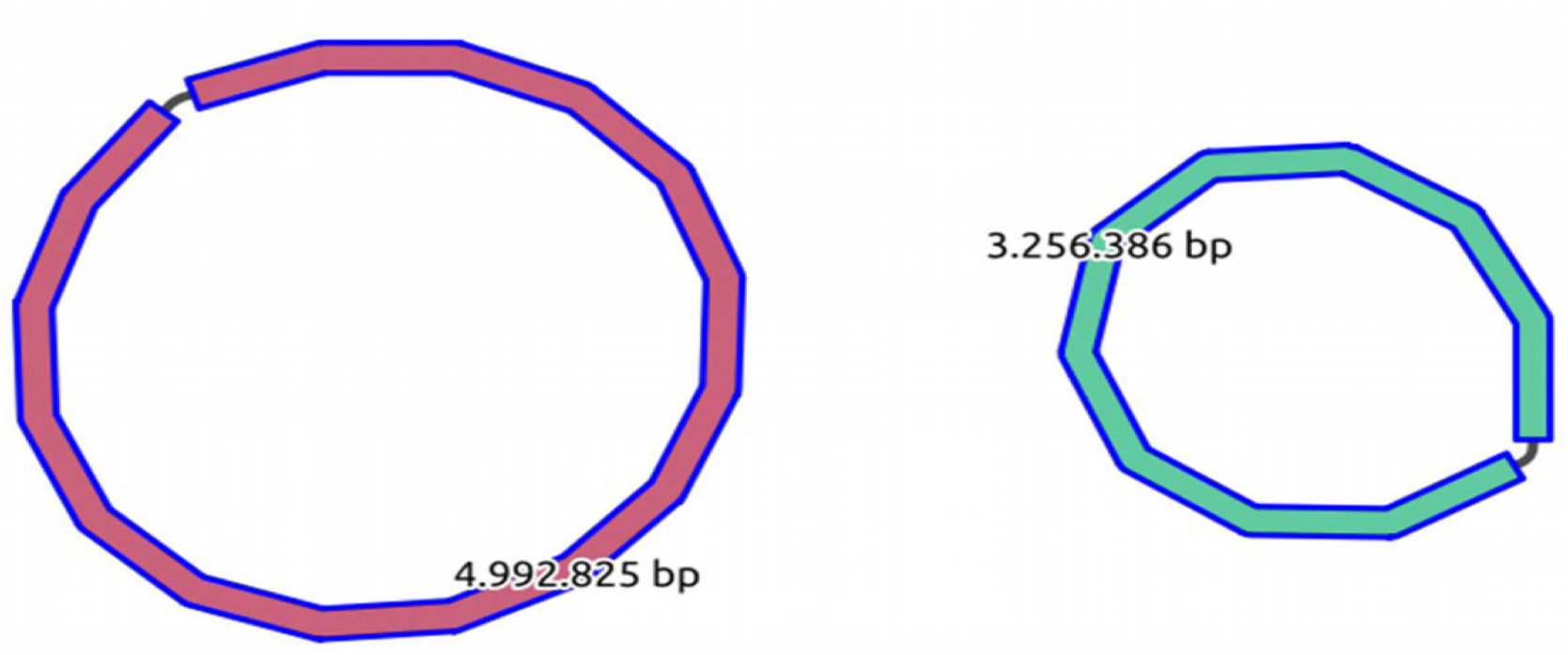
The graphical view of assembled genomes using *de novo* assembly graphs and the Bandage tool.

### 3.4. Check the Genome for Contamination

ContEst16S assisted in evaluating the assembled genomes for contamination. Including GC content, genome length, and N count, **Table 1** lists the genomic features for the smallest (3,256,386 bp) and largest (4,992,825 bp). ContEst16S matched the 16S rRNA gene segments from the assembled genomes to a database of 69,745 genomes including 596 contaminated entries. The produced genomes were confirmed to be uncontainted. While the smallest contig was tightly related to *Psychrobacter alimentarius*, ContEst16S’s maximum likelihood phylogenetic trees (**Figure 5**) revealed that the biggest contig of the genome was more similar to *Shewanella baltica* than to other 16S rDNA fragments.

**Figure 5:**
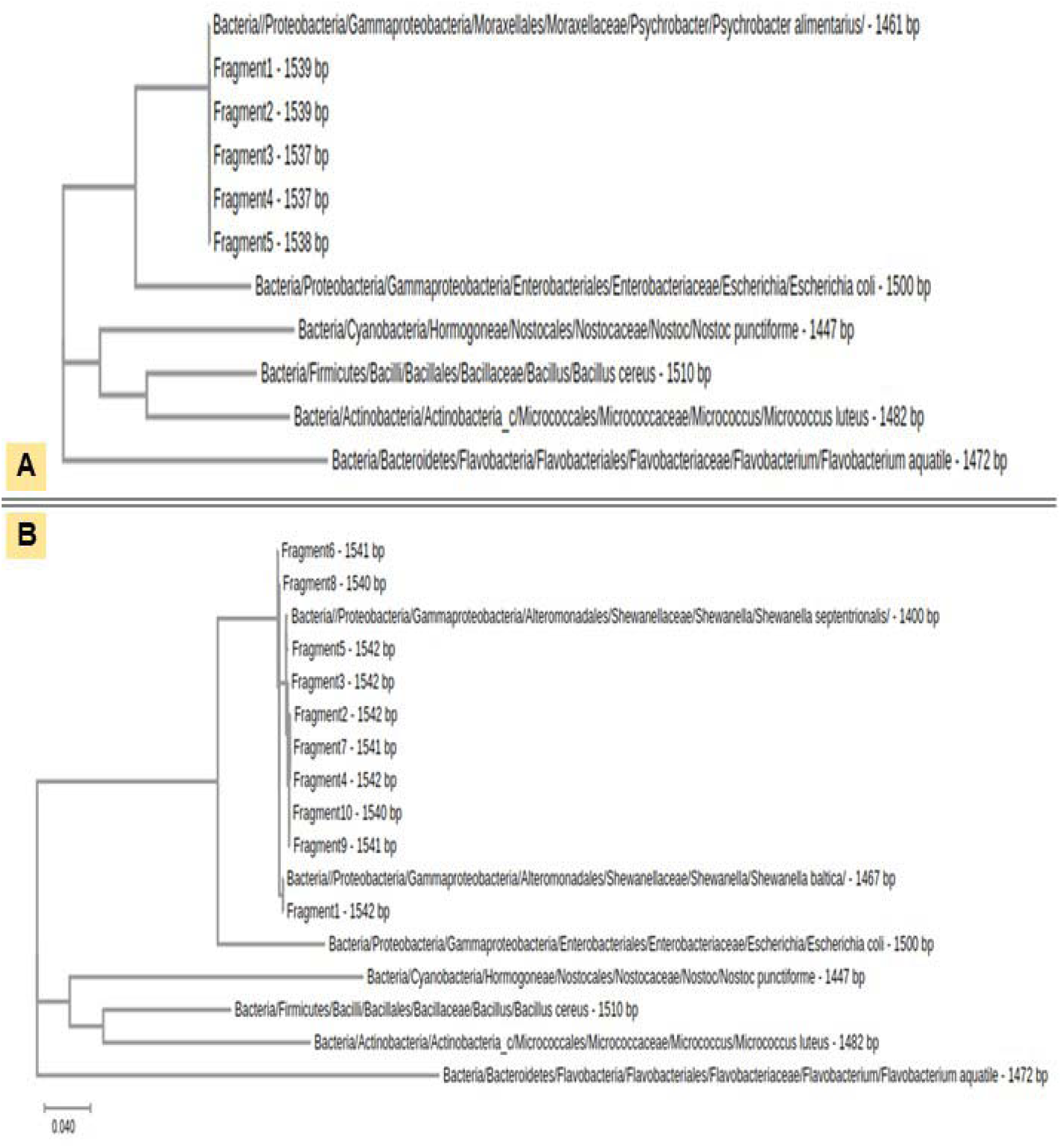
Maximum likelihood phylogenetic trees for the five fragments of 16s rDNA regions. **A**) Phylogeny of the smallest contig (3,256,386 bp). **B**) Phylogeny of the largest contig (4,992,825 bp).

**Table 1:** Features of the assembled genomes analyzed using ContEst16S.

| Contig | Total Length (bp) | A | C | G | T | N | GC Content (%) | Decision |
| --- | --- | --- | --- | --- | --- | --- | --- | --- |
| Smallest contig | 3,256,386 | 926,885 | 701,606 | 699,426 | 928,469 | 0 | 43.02 | Not Contaminated |
| Largest contig | 4,992,825 | 1,338,502 | 1,160,023 | 1,157,916 | 1,336,384 | 0 | 46.43 | Not Contaminated |

### 3.5. Fast Annotation and Submission Tool (DFAST) to DDBJ

After annotations using the DFAST tool, the two assembled genomes were turned into DDBJ’s GenBank database. Measuring 3,256,386 bp, the smallest-sized genome was annotated as *P. alimentarius* and assigned the accession number AP031495. By contrast, with 4,992,825 bp, the biggest genome was annotated as *S. baltica* and assigned the accession number AP031461.1.

### 3.6. Annotate the Complete Genome

The two assembled genomes were annotated via uploading to the server of the Bacterial and Viral Bioinformatics Resource Centre (BV-BRC) to generate the output JSON file, which was visualized using Proksee as shown in the circular map view for *P. alimentarius* and for *S. baltica* (**Figure 6**).

**Figure 6:**
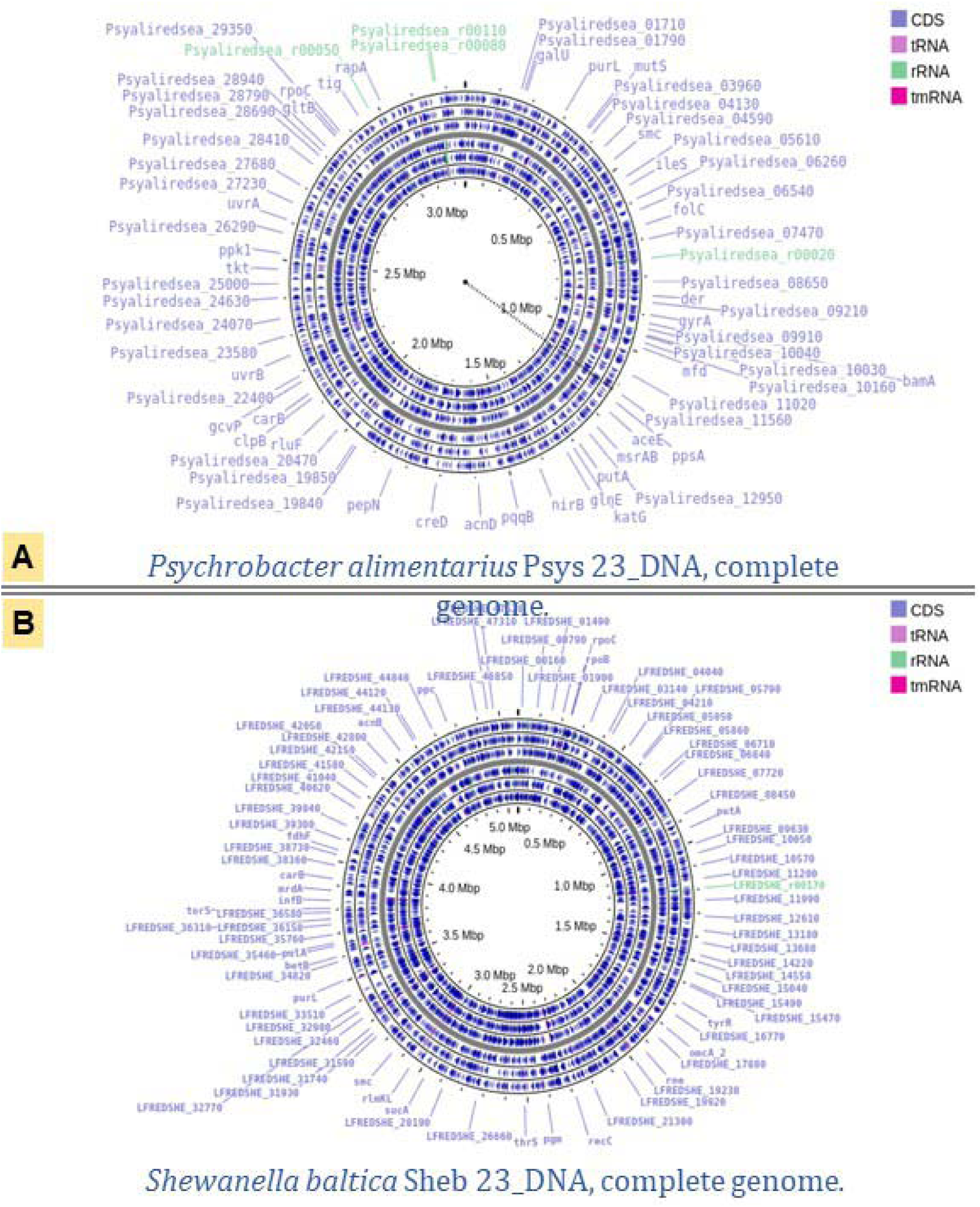
Circular maps of complete genome. **A**) Map of *P. alimentarius* (Genbank accession no. AP031495.1). **B**) Map of *S. baltica* (Genbank accession no. AP031461.1).

### 3.7 Finding the Bioluminescent Gene in the Complete Genome of *P. alimentarius*

Bioluminescent genes were identified using the PATRIC annotation tool included within the BV-BRC server. It identified the family of alkanal monooxygenase (FMN-linked) proteins by means of the GO term GO:0047646. **Table 2** offers a synopsis of the annotations, including Alkanal Monooxygenase Alpha Chain’s coding sequence (CDS).

**Table 2:** PATRIC annotation output for *P. alimentarius*.

| Feature Type | Start | End | Length (bp) | Length (AA) | Product |
| --- | --- | --- | --- | --- | --- |
| CDS | 1,193,850 | 1,194,887 | 1,038 | 345 | Alkanal Monooxygenase Alpha Chain |

### 3.8. Finding Genomes Similar to the Assembled *P. alimentarius* Bacterial Strain

To evaluate similarity between public genomes, genome distances were computed applying the Mash/MinHash technique. With an OrthoANIu value of 97.55%, the assembled *P. alimentarius* genome (AP031495) and the reference strain PAMC 27649 (CP014945) showed a high degree of similarity (**Table 3**). The assembled genome (*P. alimentarius* 2024) is tightly related to *P. alimentarius* PAMC 27889 according to a phylogenetic tree produced with the BV-BRC Bacterial Genome Tree Service (**Figure 7**). Together with accessions OR621168.1 and LK931862.1, fragments 1–5 (AP031495.1) clustered tightly to confirm that they are all bioluminescent strains (**Figure 8**).

**Figure 7:**
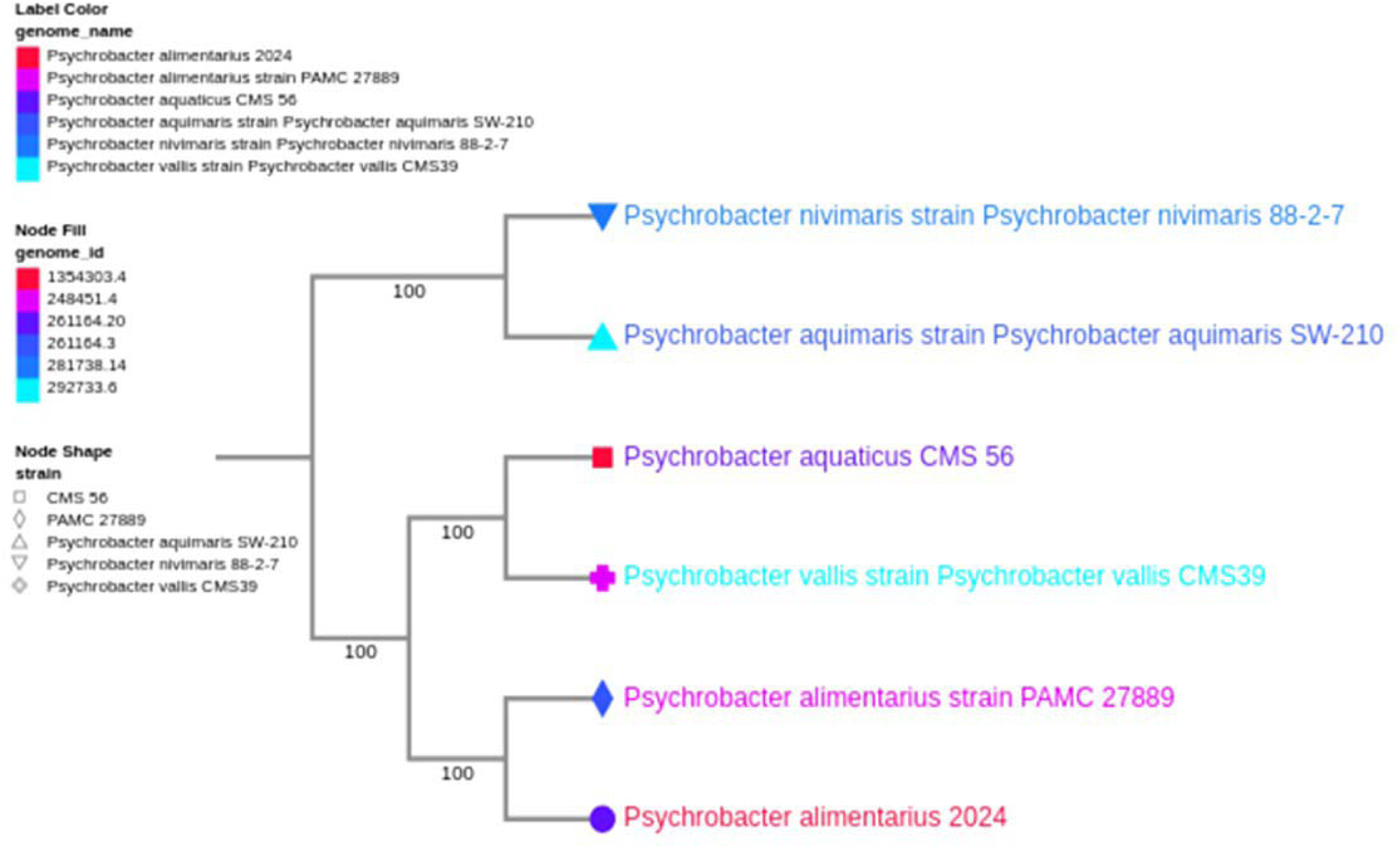
Phylogenetic links of *Psychrobacter* strains. With *P. alimentarius* strains (2024 and PAMC 27889) forming a strongly supported clade (bootstrapped >90%), The Codon Tree method picks single-copy BV-BRC reveals distinct clustering of five species. While *P. aquimaris* SW-210 shows as an evolutionary outlier with longer branch lengths, implying possible niche specialisation, moderate genetic distances separate *P. aquaticus* CMS 56 from this group. Node forms serve as strain identifiers; filled circles show strains incorporated in downstream analyses. The strong bootstrap values (>85% for all major nodes) highlight both conservation within species and divergence between them, most likely reflecting adaptations to different cold environments, so confirming the dependability of these phylogenetic relationships.

**Figure 8:**
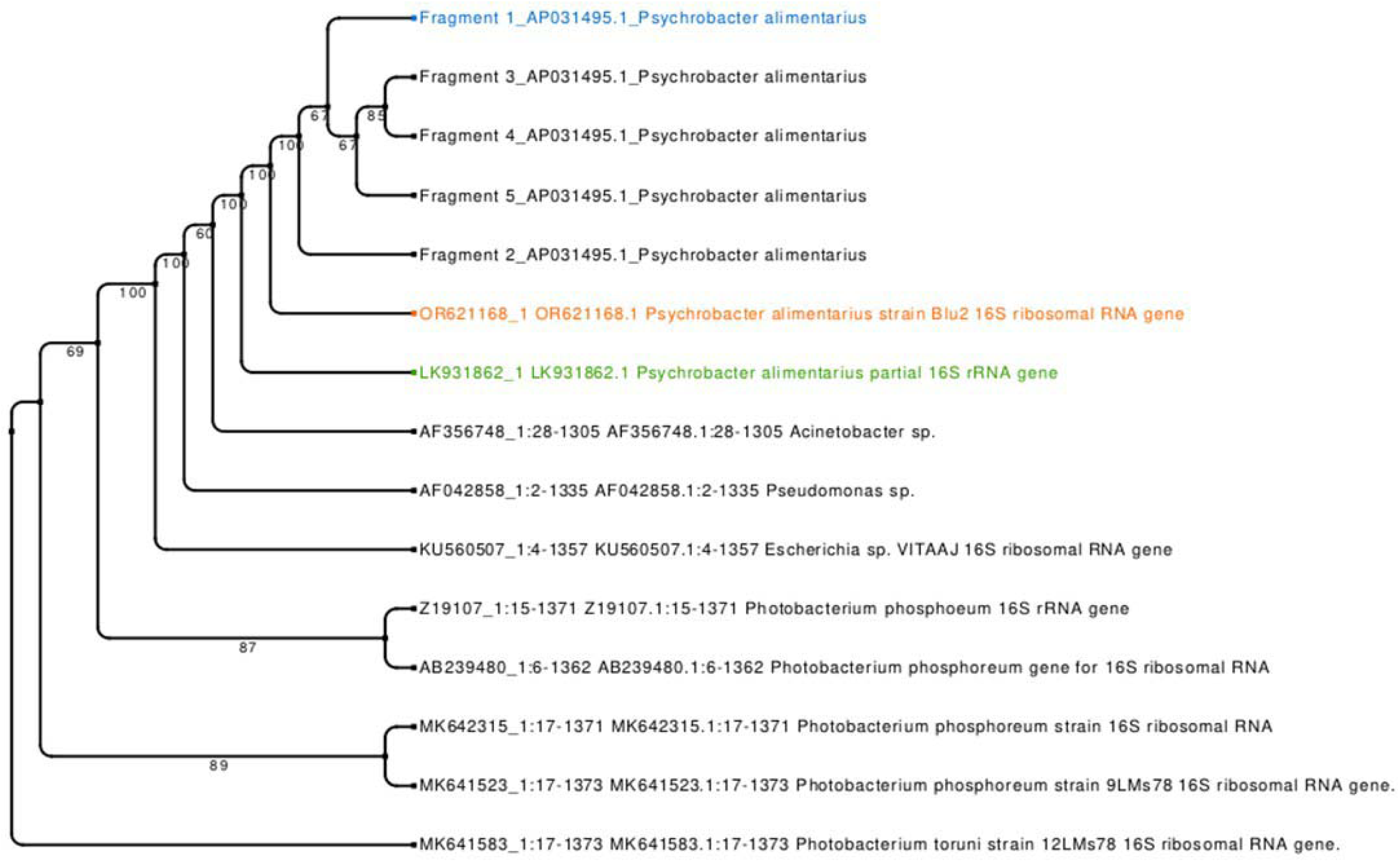
Phylogenetic tree built by MAFFT alignment and neighbor-joining of 16S rRNA sequences placed five *P. alimentarius* fragments (AP031495) in a strongly supported clade (bootstrap >95%), so verifying their taxonomic classification. Indicating great conservation, close relatives (strains Blu2 [OR621168.1] and LK931862.1) grouped within this cluster. While *Pseudomonas* sp. and *Escherichia* sp. provided outgroups, *Photobacterium* spp. developed a clear sister clade with horizontal transfer of bioluminescent genes to *Escherichia*. Branch lengths proposed higher variation among *Photobacterium* strains than in *P. alimentarius*.

**Table 3:**
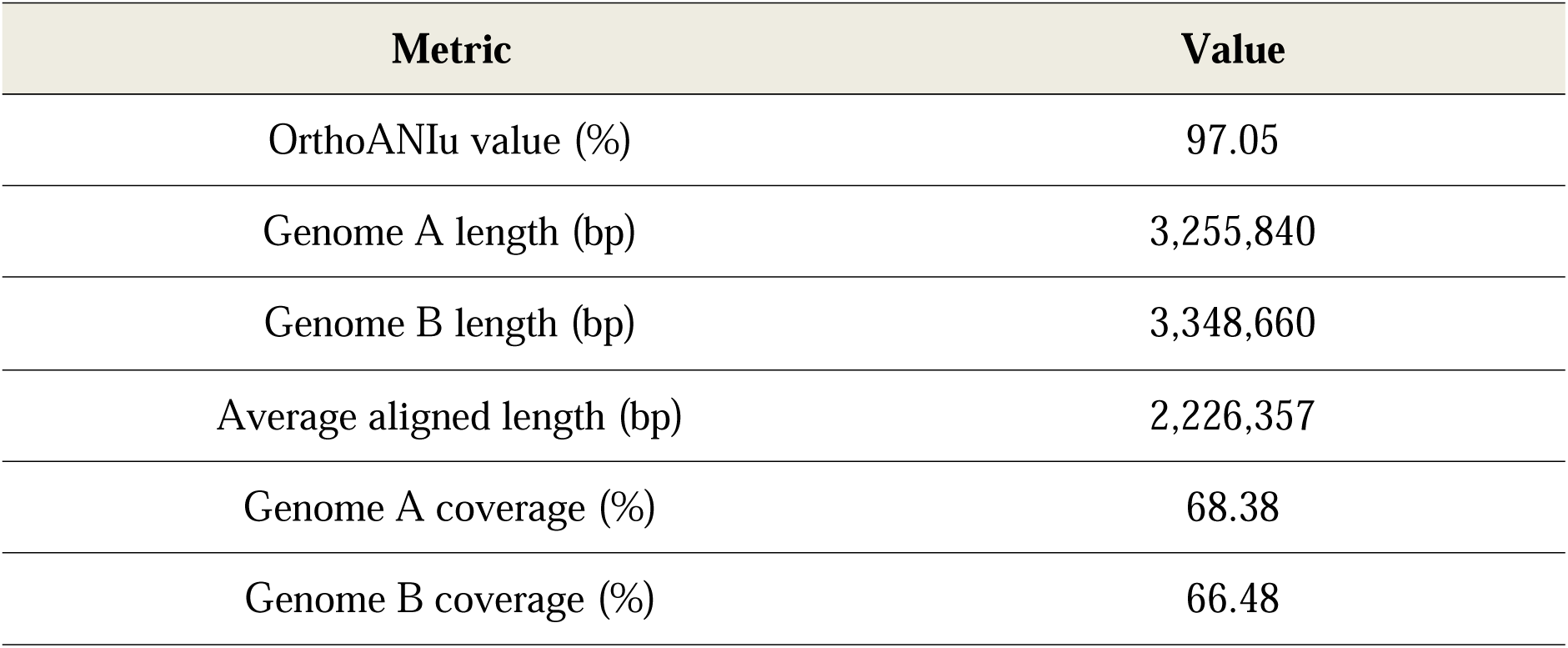
OrthoANIu comparison of *P. alimentarius* genomes.

### 3.9. Genome alignment (Mauve)

The Mauve tool brought the reference genome (CP0149) into line with the recently produced genome (AP031495.1) of *P. alimentarius*. Local collinear blocks (LCBs) found here revealed conserved genomic segments with structural integrity. **Figure 9** shows two LCB displaying homologous backbone sequences.

**Figure 9:**
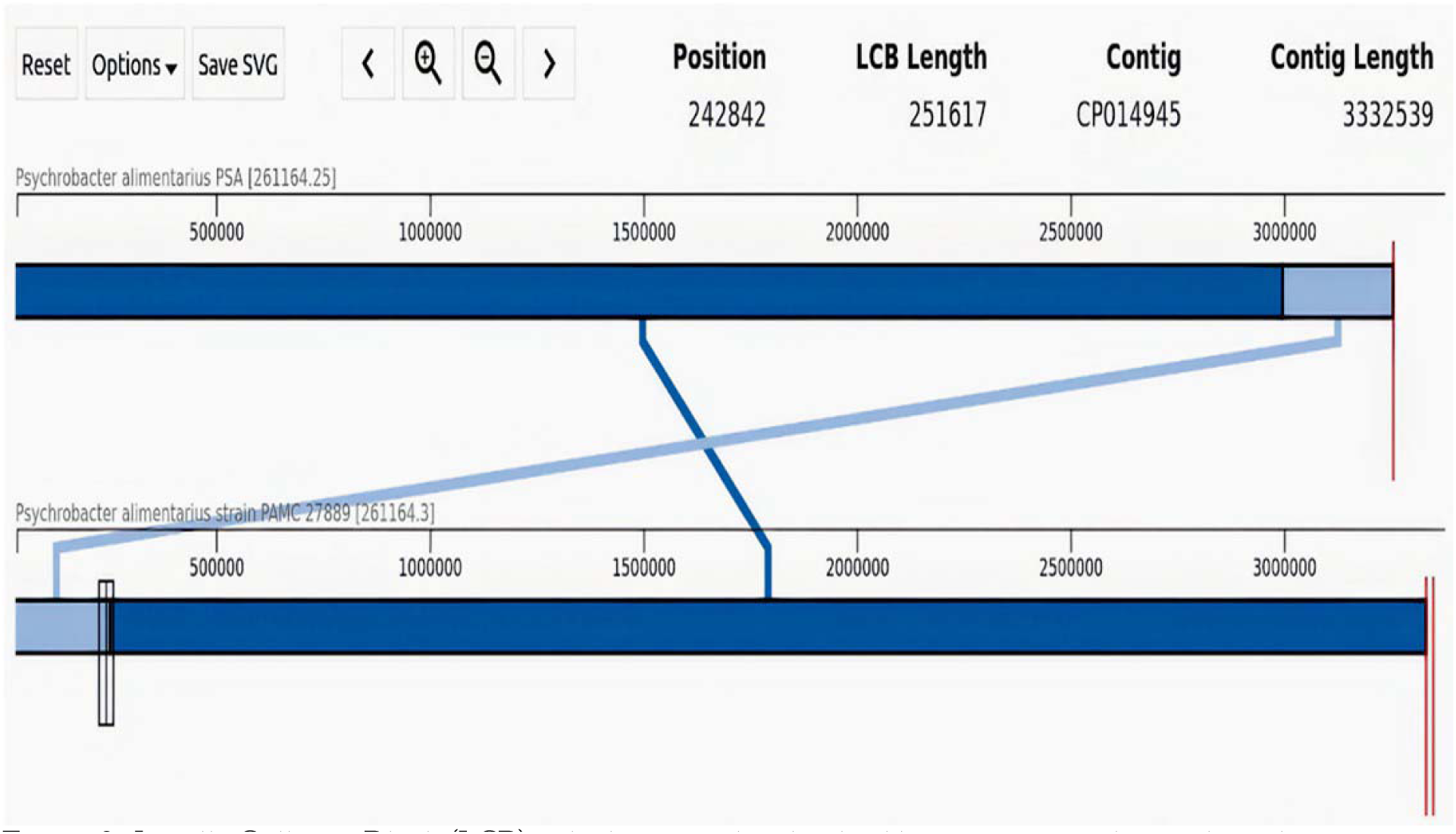
Locally Collinear Block (LCB), which is considered as backbone sequences that are homologous.

## 4. DISCUSSION

Symbiotic interactions between cephalopods and bioluminescent bacteria best illustrate the essential ecological adaptation of bioluminescence in marine life. Although the *Uroteuthis* genus is well-documented for such symbioses, their frequency throughout the Loliginidae family is yet unknown (Anderson *et al*., 2014; GenBank OR621168.). Our work separates and identifies bioluminescent bacteria from *Loligo forbesi* in the Red Sea, so offering genomic and ecological understanding of these symbiotic interactions and their evolutionary consequences.

### Important discoveries and genomic perspectives

We discovered two bacterial symbionts in *L. forbesi* using Nanopore sequencing: *S. baltica* and *P. alimentarius*. The effectiveness of long-read sequencing for resolving complex microbial genomes was highlighted by the discovery of few dark regions in high-quality genome assemblies (Ebbert *et al*., 2019). Our isolates and reference strains (*P. alimentarius* CP014945.1, *S. baltica* CP002384.1) were found to have close relationships, according to phylogenetic analysis. Locally collinear blocks (LCBs) highlighted conserved genomic architecture. In line with earlier reports of low-light emission in this genus, *P. alimentarius* was notable for having a LuxA domain associated with dim bioluminescence ([15] Isolation, identification, and manipulation ). By identifying bioluminescent *P. alimentarius* in *L. forbesi* for the first time, its known associations within Loliginidae are expanded.

### Ecological and evolutionary implications

*Shewanella* and *Psychrobacter* have clearly established mutualistic roles in marine symbioses. While Psychrobacter adds to host fitness by oxidative stress reduction, *Shewanella* spp. guard squid eggs from pathogens and abiotic stress (Barbieri *et al*., 1997). Our identification of bioluminescence-related genes in *P. alimentarius*—including the alkanal monooxygenase family (GO:0047)—indices adaptive advantages in dimly lit environments, such UV protection or prey attraction. These results support current studies on bioluminescent *P. alimentarius* in *Loligo vulgaris* (GenBank OR6211), so highlighting the ecological adaptability of the genus.

### Bioluminescence as a Trait Under Conservation

Fascinatingly, non-luminous bacteria such as *Vibrio cholerae* retained truncated *lux* operons, generating barely detectable light to humans. This is consistent with observations in *Enhygromyxa salina* whereby low-intensity luminescence (Yudenko *et al*., 2024) results from a simplified *lux*ACDE operon. Even in species with functional expression lost, such “cryptic” bioluminescence suggests evolutionary conservation of light-production machinery (Ramaiah *et al*., 2000). This phenomena can be explained by regulatory disturbances, incomplete lux paths, or ecological niche shifts; hence, research on gene expression dynamics is justified.

### Future Directions

Our work emphasizes the need of investigating bioluminescence control among bacterial taxa and surroundings. Comparative research of *lux* operon organization, transcriptional control, and ecological pressures will help to define its evolutionary path. Furthermore, underdeveloped are the uses of bioluminescent bacteria in environmental monitoring, including biofilm indicators or pollution biosensors.

## 5. CONCLUSION

Finally, this work reveals genomic preservation in *Psychrobacter* and cryptic *lux* genes in non-luminous taxa, so advancing knowledge of bioluminescent symbioses in cephalopods. We highlight the dynamic adaptive function of bioluminescence in marine ecosystems by linking genomic, ecological, and evolutionary points of view.

