## Supplementary figures and images for "Nanopore-Based Whole-Genome Sequencing Reveals a Bioluminescent and Non-Bioluminescent Bacterium Isolated from the Squid *Loligo forbesi* in the Red Sea"

### Supplemental Data 1

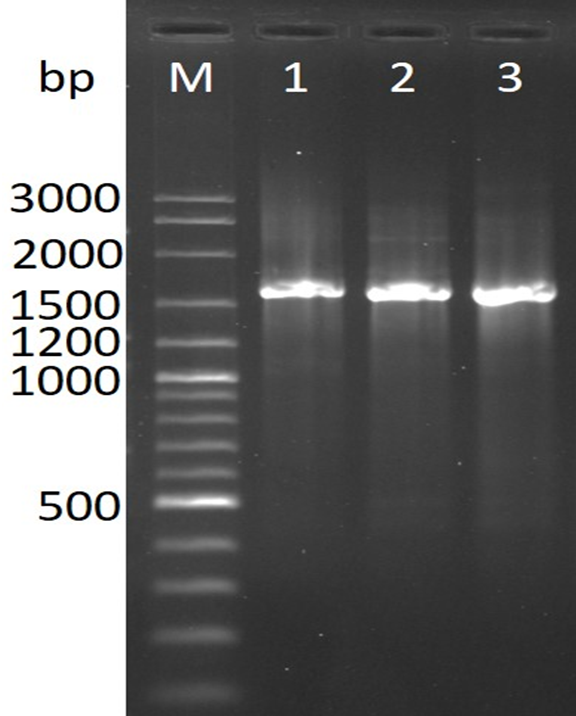
